# Traumatic Conditioning Induces a Combination of Anesthesia Resistant Memory and Protein Synthesis Dependent Long-Term Memory in Adult *Drosophila*

**DOI:** 10.64898/2025.12.07.692887

**Authors:** Snehasis Majumder, Gaurav Das, Abhijit Das

**Author notes:** Corresponding authors: Abhijit Das, Gaurav Das.

## Abstract

Stressful experiences elicit long-lasting memory trace that often persists for life-time. Though storage of experience dependent memory is key to survival, post-traumatic memories are detrimental to the physical and emotional wellbeing. The generation of any form of long-term memory culminates from synaptic plasticity and potentiation through persistent change in protein synthesis. *Drosophila* has been a robust model for the study of memory mechanisms; olfactory memory paradigms have been enormously successful in elucidating neuroanatomical, molecular, physiological and signalling pathways underlying the process. Long-term memory (LTM) has been induced by protein synthesis dependent mechanisms followed by a post-learning consolidation process whereas a medium-term anaesthesia resistant memory (ARM) is mediated by the Radish protein in flies. Here we present a novel and simple memory paradigm in adult flies for inducing sustained stress-induced memory paradigm, where a single attractive odorant is associated with an aversive stimulus-copper sulfate, which provides bitter taste as well as sustained malaise due to toxicity. We find that the elicited memory is a combination of ARM and LTM which is dependent on the number of training cycles. The eight-cycle training paradigm leads to robust, CREB and protein synthesis dependent memory persisting for about 7 days. The mushroom body (MB) Kenyon cells neurons as well as the dopaminergic input neurons (PPL1 subset) are involved in the formation of memory mediated by enhanced calcium as well as synaptic elaboration in the γ subset of the MB neurons.

## Introduction

Encoding of experiences in the form of memory and their future retrieval is fundamental for the survival of any organism. It is an instinctive survival drive to perceive and adapt to the dynamic surroundings by bringing about changes in the respective neuronal engrams, leading to behavioural plasticity. Conversely, traumatic experiences generate long-term memory that sometimes persists for lifetime and culminates in pathological stress, anxiety and depression. It is of great significance to understand the mechanistic basis of the formation of traumatic memory, which originates from sustained adverse experience. Olfactory learning in the model organism *Drosophila* has been instrumental in understanding the fundamental molecular, physiological and synaptic mechanisms governing the process. Decades of research using mutant and transgenic flies as well as pharmacological treatment have led to the classification of olfactory avoidance memory into four distinct types, including short-term memory (STM), middle-term memory (MTM), anaesthesia-resistant memory (ARM), and long-term memory (LTM) (Tully et al.; Margulies et al. 2005). STM persists in the fly for a very brief period of up to 60 minutes after conditioning. Genetic mutants like *dunce* (a cAMP phosphodiesterase mutant) and *rutabaga* (a Ca^2+^/calmodulin activated adenylyl cyclase mutant) first introduced severe STM impairment, underscoring their role in the protein synthesis independent (PSI) cAMP signalling pathway for STM formation, which is consistent with earlier discovery in the *Aplysia* memory model (Davis and Kiger 1981; Livingstone et al. 1994; Tully and Quinn 1985; Schwartz et al. 1971; Kandel 1995). The intermediate medium-term memory (MTM) is formed after STM and prior to ARM (Margulies et al. 2005). Mutations in cAMP-dependent protein kinase A (PKA), *DC0,* showed disrupted MTM retention, which was identical to the primarily discovered *amnesiac* mutants (Li et al. 1996; Skoulakis et al. 1993; Tully and Quinn 1985; Tully et al. 1990). ARM is a unique memory type in the flies that is not susceptible to hypothermic pulse. Mutations in *radish* gene, which codes for RAP like GTPase activating protein, disrupts ARM (Tully et al. 1994), but the LTM remains unaltered. Both ARM and LTM were known to arise simultaneously in flies, even though a recent arguable hypothesis indicates that the ARM and LTM formation is mutually exclusive and solely depends on either massed or spaced training (Tully et al. 1994; Isabel et al. 2004). LTM lasts longer than ARM and demands for de novo expression of new genes, hence, protein synthesis dependent (PSD) (Tully et al. 1994). In contrast, STM, MTM and ARM are independent of synthesis of new proteins. Transcription of genes like *CrebB*, *nalyot*, *notch*, *nebula*, *pumilio, crammer* etc. are required for the induction of LTM (Yin et al. 1994; James DeZazzo et al. 2000; Presente et al. 2003; Chang et al. 2003; Josh Dubnau et al. 2003; Comas et al. 2004).

Classical conditioning, which establishes correlations between simultaneous events, is the basis of most of the learned behaviors of organisms including humans. The classical conditioning memory has been studied in multiple organisms including *Drosophila* (Tully and Quinn 1985). The olfactory conditioning paradigm in the fruit flies is a powerful experimental tool to decipher anatomical, molecular and physiological pathways contributing to memory formation (Tully and Quinn 1985). The classic paradigm entails association of an attractive odorant with a punishing experience. The flies are exposed to two attractive odorants consecutively, one of the odorants (conditioned stimulus, CS+) is coupled with an electric shock (unconditioned stimulus, UCS), followed by solo presentation of a second odorant (CS-); training the flies by repeated presentation of this combination of the ‘CS+ with UCS’ and the ‘CS-alone’ leads to formation of a strong aversive memory in the flies towards the CS+ odorant (Tully and Quinn 1985). In addition to electric shock as a UCS, the bitter taste of several compounds like quinine and caffeine is also applied to elicit ‘odor-bitter taste’ coupling memory (El-Keredy et al. 2012; Apostolopoulou et al. 2016). In an aversive conditioning paradigm (Mohandasan et al. 2022), it has recently been observed that flies formed long-term aversive associative memory when an attractive odorant was coupled with the bitter taste of CuSO_4_.

For *Drosophila*, the mushroom body (MB) receives multimodal sensory inputs for processing and integration into a complex behavioral outcome. Moreover, MB neurons display experience-dependent synaptic plasticity leading to behavioral plasticity. MB acts as an associative learning center in the protocerebrum; enabling flies to establish connections between the odor (conditional stimulus) with rewards/punishments (unconditional stimulus) to form either appetitive or aversive memories respectively (Aso et al. 2014). To enable this association, two separate clusters of dopaminergic neurons (DANs) project their axons within different lobes of MB and transmit either reward or punishment sensation to specific regions of MB. In the appetitive learning, the rewarding input is conveyed to the KCs-MBONs (Kenyon cells-Mushroom body output neurons) by DANs of the PAM (Protocerebral anterior medial) cluster. Further, in case of aversive learning, posterior DANs of PPL1 or PPL2 (Protocerebral Posterior lateral) cluster relays negative reinforcement onto the KCs-MBONs connection (Aso et al. 2014). Though the odor-shock coupling based aversive conditioning has elucidated the fundamental molecular, signaling and transcriptional pathways underlying long-term memory, the memory induced through such training is ‘acute’ in nature and would not qualify as a model for sustained traumatic memory. A ‘chronic’ or sustained adverse experience coupled with an odorant in repeated manner through spaced training is required.

Here we present a novel and simplified aversive olfactory conditioning paradigm where a bitter and mildly toxic CuSO_4_ feeding has been used as UCS to replace the electric shock and single odor as CS+, i.e., excluding the necessity of CS-odorant from the paradigm. CuSO_4_ provides two ways to generate sustained adverse experience in flies: through bitter gustatory experience (Xiao et al. 2022) and toxicity-based malaise (Balinski and Woodruff 2017; Halmenschelager and da Rocha 2019). Neuronal correlates of this memory paradigm, as well as the requirement of CrebB, have been elucidated. Additionally, this newly optimized memory paradigm has already proved its potential to address learning and memory impairment associated with Tauopathy and its genetic rescue (Bisht et al. 2024). Here, we used this paradigm to shed light on the long-standing questions regarding the ARM and LTM; we show that STM, ARM and LTM forms in training cycle dependent manner, and rather than being exclusive, both ARM and LTM are formed simultaneously using distinct molecular pathways, contributing to the aggregate memory available at any given time. We establish the contribution of both MB neurons and punishment-specific dopaminergic neurons (PPL1) neurons to be involved in the memory formation. Role of punishment-specific dopaminergic neurons (PPL1) during consolidation suggests the persistence of post-training reinforcement even without external cue in case of long-traumatic memory.

## Materials and Methods

### Fly Stocks

*Canton S* (wild type), *UAS mCD8GFP (Bl-5137), UAS TRIC (Bl-62827), UAS CrebB RNAi (Bl-63681),* lines were obtained from the Bloomington Stock Centre, Indiana, USA. *Radish* mutant (*rsh^MI12368-TG4.1^; Bl-76737), MB247-Gal4* (Bl-50742)*, UAS Shibire (ts1)* (Kitamoto 2001)*, MB504B-Gal4* (Aso et al. 2014b), were generously gifted by Dr Gaurav Das from NCCS Pune.

### Fly maintenance

Flies were reared in standard cornmeal media in plastic bottles in 25°C incubators with a 12-hr light/dark (LD) cycle.

### Training paradigm for aversive stress conditioning in adult *Drosophila*

For the induction of aversive LTM in adult *Drosophila*, the following procedure was used (Figure 1A):

1) Following three types of media vials were prepared:
  a. Starvation media containing 0.75% agar.
  b. Control media contained 0.75% agar with 85 mM sucrose.
  c. Punishment media was prepared by mixing freshly autoclaved control media with 80 mM CuSO_4_ at around 55℃-60 ℃. The media is then constantly agitated for one minute to dissolve all the Copper Sulphate crystals.
2) About 5 ml of each media was gently poured into the designated training vials (borosilicate glass, outer diameter 25 mm and height 85 mm) without forming bubbles.
3) 1000 µl of 5% (v/v) 2,3 Butanedione (2,3 BD; attractive odorant) was prepared with autoclaved double distilled water. A 2 cm X 1.5 cm filter paper was soaked with 100 µl of 5% (v/v) 2,3 BD. Each odor-soaked filter paper was placed in a 3D-printed odor cup specially designed for delivering odor into the respective test tubes. Eight such odor cups were used to perform one round of experiment: four for the control group while the other four for the punishment group.
4) 4/5 days old flies (both sexes) were collected and kept in freshly prepared food for 24 hours.
5) Then they were starved in starvation media for 12/16 hours before training.
6) Prior to training, both the control and punishment vials were odorized with 2,3 BD for at least 20 minutes.
7) Each odorized control and punishment vial were used for training not more than two groups of flies.
8) The ‘naïve group’ flies were exposed to 100 µl of 5% 2,3 BD through soaked filter paper in the control media vials for 5 minutes followed by 5 minutes of incubation in an empty test tube. This 10-minute training was considered as one naïve training cycle.
9) The training for aversive conditioning was performed by exposing the ‘trained group’ of flies to 100 µl of 5% 2,3 BD in punishment media vials for 5 minutes followed by 5 minutes of incubation in an empty test tube. This 10-minute training was considered as one punishment cycle.
10) Eight such training cycles were performed for both naïve and punishment groups.
11) To check memory retention 12 hrs post-training, the flies were starved in starvation media for 12 hours followed by Y maze assay. For checking one-day memory retention, flies were starved 6 hours after training followed by a 5-minute food pulse; these flies were again starved for 18 hours prior to Y-maze assay. For checking memory retention beyond 1 day, trained or naïve flies were starved for 6 hrs and were then shifted to normal food. Afterwards, the flies were again starved for 14 to 16 hours prior to Y maze assay.

**Figure 1.**
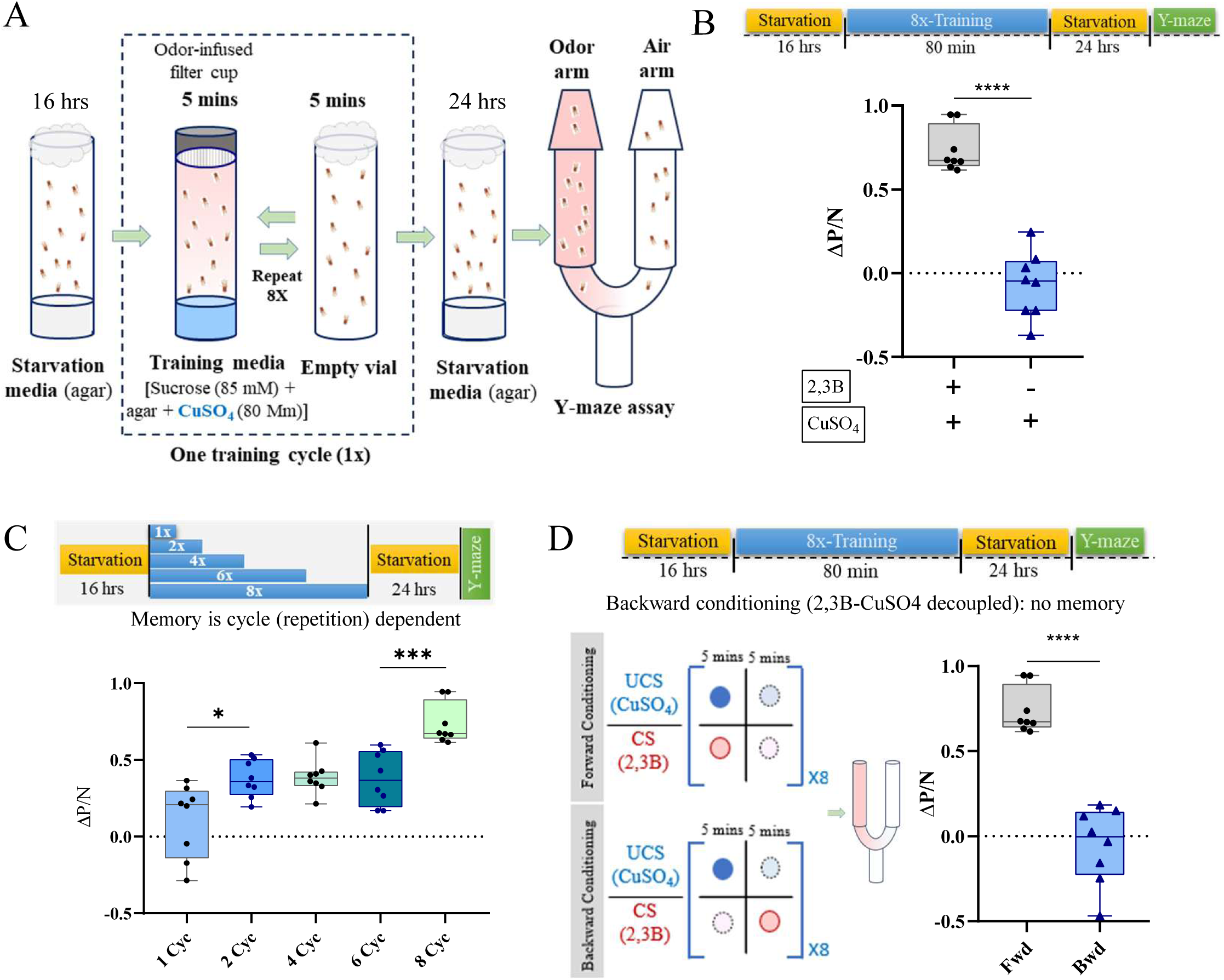
Association of an attractive odorant with bitter/toxic Copper Sulphate (CuSO_4_) in food elicits strong long-term aversive memory in adult flies. A. Schematic diagram illustrating the training and testing paradigm of “odor-bitter food” associative conditioning in adult flies eliciting aversive long-term memory. Flies were subjected to 16 hrs starvation followed by 8X training cycles (unless otherwise specified), each cycle consisting of 5 minutes in a vial containing CuSO_4_ in sucrose-agar media followed by 5 minutes in an empty vial. The memory scores (Performance Index) were obtained 24 hrs after conditioning in a Y-maze assay (see methods section for full description). [The gap between training and Y-maze assay were changed to suit the need of individual experiments, the same has been indicated on top of each figure panel in all subsequent figures]. B. After one day post-memory induction, the ΔP/N value was significantly increased for the trained group of flies compared to the ‘no-odorant’ control group, indicative of formation of associative memory. C. Wild-type flies were trained with variable number of training cycles (1 Cyc, 2 Cyc, 4 Cyc, 6 Cyc and 8 Cyc) and their 24 hrs memory was assessed. Flies trained with one training cycle (1 Cyc) retained their memory index, ΔP/N value closer to zero, suggesting no memory retention. 2 Cyc-trained flies exhibited a significant increase in ΔP/N value compared to 1 Cyc trained flies, indicating training induced memory formation. 8 Cyc trained flies also showed a similar significant increase of ΔP/N value with respect to 6 Cyc trained flies after 24 hrs. D. Forward conditioning showed normal memory formation (same data points as Figure 1B) whereas memory performance of flies trained with *backward conditioning* where CuSO_4_ and odor were presented in decoupled fashion (CuSO_4_ in one vial and odor in another vial) and tested after 24 hrs revealed significant drop in ΔP/N value compared to forwardly trained group; this shows no sign of LTM after backward conditioning. N=8 biological replicates in each behavioral experiment. Bars represent the Mean ± Standard Error of Mean (SEM). Unpaired two-tailed t test was performed for Figure 1B & Figure 1D. One-way ANOVA followed by Tukey’s multiple comparison test was performed for Figure 1C. **** signifies p value < 0.0001, *** signifies p value < 0.001, ** signifies p value < 0.01, * signifies p value < 0.05, ns (non-significant) signifies p value > 0.05.

### Testing of olfactory memory after induction of LTM in adult flies

A classic glass Y-maze apparatus was used to test the odor choice in the naïve and the trained group of flies. The Y-maze assay set-up and the method were following (Das et al. 2011; Dey et al. 2024). The glass apparatus resembles the shape of the letter ‘Y’ and was positioned upright for the assay. The odorant (2,3 BD) was continuously passed through one of the arms by bubbling air through a solution of 10^-3^ (v/v) 2,3-BD in a bottle and pumping it through the ‘odor arm’. Air bubbling through distilled water was continuously pumped through the ‘air arm’ of the Y-maze. The flies were placed in the entry tube on the bottom of the maze and due to the negative geotaxis behavior of the flies they try to climb upward, enter the Y-maze and choose to enter the odor arm or the air arm according to their preference (Twick et al. 2014). Number of flies in each of the arms were counted after one minute and a performance index was calculated to represent their attraction towards the odorant based on the following formula:

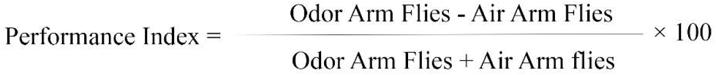

After calculating the Performance Index (PI) of naïve and trained flies the mathematical readout of the memory ΔP/N is calculated by the following formulae.

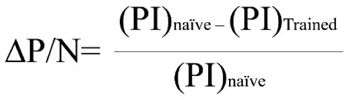

ΔP/N values of the respective groups were compared statistically using GraphPad Prism software.

### Induction of anesthesia in adult flies by cold exposure to check anesthesia resistant memory (ARM)

Post-training flies were exposed to ice cold temperature to induce anesthesia. First, folded tissue papers were inserted into empty glass vials to absorb extra moisture during cold exposure. These vials were then pre-cooled by submerging them up to their cotton plugs in an ice tray for 5 minutes. The naïve and trained flies were transferred into separate pre-chilled vials for cold shock for 2 minutes. Flies typically lose consciousness approximately 30 seconds into the cold exposure and begin to regain consciousness around 30 seconds after removal from the cold. Following cold shock, flies were transferred to starvation media for 6 hours, after which they received a brief 5-minute food pulse. This was followed by an additional 18-hour starvation period prior to performing the Y-maze behavioral assay to assess 1-day memory retention. For checking impact of anesthesia on 3h memory, cold-shocked flies were starved for entire 3h followed by Y-maze assay.

### Blocking neurotransmitter release from neurons by *Shi^ts^*

For blocking the release of neurotransmitter from neuronal subsets, temperature sensitive dynamin (*Shibire^ts^*) was expressed. The *Shi^ts^* proteins were activated by exposing the adult flies to the higher temperature (30℃ – 32℃) (Kitamoto 2001). For blocking neurotransmission during training, the 18℃-reared overnight starved flies were transferred to the elevated temperature (32℃) 30 minutes before the memory induction and the entire training was conducted at 32℃. Post-training, flies were returned to starvation media at 18℃ with a short food pulse of 5 minutes in between after 6 hours until the 1 day memory testing by Y-maze assay. For blocking neurotransmission during consolidation, training was performed at 18℃ and post-training, the flies are transferred to 32℃ for 6 hours in starvation media followed by 5 min food pulse and 18 hours in 18℃. The memory score was obtained after 1 day through Y-maze experiment performed in room temperature.

### Cycloheximide treatment for blocking protein synthesis

To reduce protein synthesis in the brains of adult *Drosophila* during conditioning, wild-type flies are fed with 35 mM Cycloheximide (SRL #86620) before the training session as described in (Tully et al. 1994). Cycloheximide 0.75% (w/v) was added to agar media after autoclaving at ∼ 60℃. Wild-type flies were allowed to feed on the freshly prepared Cycloheximide-Agar media for 16-18 hours prior to conditioning to ensure adequate ingestion of the drug.

### Validation of CrebB knockdown using CrebB-shRNA

To validate the knockdown of CrebB, semi-quantitative method was used. *UAS-CrebB RNAi* was crossed to *Tub-GAL4* to express the RNAi construct in all cells. Adult fly heads of the wild type flies were used as a control. 10 adult heads of each genotype were homogenized, mRNA was extracted using RNeasy mini kit (#74104, Qiagen), cDNA was synthesized using Verso cDNA synthesis kit (#AB1453A, Thermo Scientific) and the *CrebB* cDNA was PCR amplified using *CrebB* gene-specific primers (forward primer: AGAATGCGGCCGCTAATGGACAACAGCATCGTC and reverse primer: ACCGACTAGTCTAGATCAATCGTTCTTGGTCTGACAG). PCR products were run on 1% agarose gel to check the DNA band of *CrebB*.

### Brain dissection, Immunohistochemistry and imaging

Adult fly brains were dissected in PBS (phosphate buffer saline) and were fixed in 4% paraformaldehyde solution (in PBS) for 40 minutes. The brains were washed with PBT (PBS containing 0.1 % Triton X-100) solution, blocked in 0.1% bovine serum albumin (BSA) in PBT and incubated with primary antibody mixture for 2 nights at 4℃. Anti-Brp (1:100, DSHB #nc82-c), anti-GFP polyclonal (1:1000, ThermoFisher #A10262), Anti–Dlg (1:100, DSHB #4F3), Anti-RFP (1:100, ThermoFisher #R10367) were used as primary antibodies. After 2 overnights, again the brains were washed in PBT, incubated with secondary antibody mix for 2 hours in room temperature with gentle rotation. Secondary antibodies Goat-Anti-Chicken-FITC (Bio Legends #410802), Alexa Fluor 647 Goat-anti-mouse (ThermoFisher #A-21235), Alexa Fluor 568 Goat-anti-mouse (ThermoFisher #A-11004), Alexa Flour 568 Goat –Anti-rabbit (ThermoFisher #A-11011) were used at 1:400 dilution. After PBT wash the brains were mounted with 70% glycerol and imaged in Olympus FV3000 Confocal microscope. Images were analyzed using Fiji (ImageJ) software.

### Quantification of fluorescence signal from desired brain regions

The confocal sections containing the MB γ-lobe regions were identified and z-projection was taken from the same number of optical sections in Fiji (ImageJ) software. The desired regions of the MB lobes were selected based on the respective staining and by using the manual ROI selection tool of Fiji. The Mean Fluorescence Intensity (FI) was calculated from the z-projections of each MB. The final FI was obtained by subtracting the background intensity of the unstained region from the Mean FI of the respective stacks. The final FI values of the naïve and trained flies were compared statistically using unpaired two-tailed t-test, using GraphPad Prism software.

## Results

### An ‘attractive odorant-toxic food’ associative training paradigm triggers the formation of long-term aversive memory in the adult fly

A single attractive odorant 5% 2,3 Butanedione (CS+) and 85mM CuSO_4_ (UCS) in sucrose agar media are presented to ∼40 flies in a ‘training vial’ for 5 minutes, followed by transfer of the flies to an empty ‘incubation vial’ again for 5 minutes; this 10 minute ‘training and incubation’ cycle was repeated 8 times (Figure 1A). Post-training flies were incubated in starvation media (0.75% Agar media) for 24 hrs with 5-minute food-pulse after 6 hrs. Associative memory of the flies were assessed in a classical binary odor-choice assay in a Y-maze where flies had to make a choice between the CS+ odorant (2,3B) and air in the two arms of the Y-maze. The naїve flies, which experienced same training except they were exposed to 2,3B without CuSO_4_ in the Sucrose-Agar media, were attracted towards the odor as evident from movement of ∼80% of the flies to the odor-arm of the Y-maze. In contrast, the trained flies which had undergone 2,3B-CuSO_4_ conditioning training, have reduced attraction towards the conditioned odorant as evident from low performance index (Supplementary Figure S1A); this results in positive memory index (ΔP/N, Figure 1B), indicating strong memory formation. Another group of flies were exposed to CuSO_4_ alone without odorant coupling, these flies showed preference towards the odorant arm in Y-maze, i.e., performance index was indistinguishable from that showed by the naïve flies (Supplementary Figure S1A), resulting in the ΔP/N value near zero (Figure 1B), i.e. no memory formation without coupling between 2,3B and CuSO_4_.

### The aversive LTM is specific towards the conditioned training odorant

Subsequently, to check the specificity of memory response towards the training odorant (CS+), flies were trained to associate bitter taste with attractive odorant 4-Methylcyclohexanol (CS+) followed by a test of their odor preference with the odorant 2,3B (novel odorant). In the binary odor choice assay the flies did not show any reduction in attraction towards 2,3B, i.e. performance index of MCH-CuSO_4_ trained flies towards 2,3B was not significantly different from that of the naïve flies (Supplementary Figure S1B), indicating that aversive olfactory memory is formed only against the odorant that has been associated with the bitter/toxic food.

### The strength of the aversive conditioning memory is dependent on the number of training cycles

Since the training of 8 cycles generated a strong memory, we next aimed to understand the dependence of the training paradigm on the number of repetitions, i.e., number of training cycles. To achieve this, different number of training cycles were performed in different group of flies: 1 cycle, 2 cycles, 4 cycles, 6 cycles, followed by Y-maze test after 24 hours; the resulting ΔP/N values were compared across the training cycle groups respectively. Performance index of the flies trained with 1 cycle is not significantly different from that of the naïve flies (Supplementary Figure S1C), hence the memory index (ΔP/N) was near zero (Figure 1C), indicating lack of memory after 24 hours. In contrast, memory indices of 24 hrs memory for 2, 4 and 6 cycle trained fly groups were positive and significantly higher than 1 cycle trained flies. Moreover, the ΔP/N of 8-cycle trained flies showed a sharp increase from the ΔP/N of 6 cycles (Figure 1C). In summary, 8-cycle training was found to generate a strong and robust long-term memory, and hence the 8-cycle training paradigm was chosen for subsequent experiments.

### Conditioning is contingent upon coupling of attractive odor with the bitter/toxic food

To further establish the role of ‘associative learning’ in the aversive memory formation in this paradigm, flies were trained with backward conditioning, where, instead of presenting the unconditioned stimulus (bitter/toxic CuSO_4_) and conditioned stimulus (odorant 2,3B) together, they were presented in successive manner. The flies were presented with CuSO_4_ in the training vial without the odorant followed by the exposure to the conditioned stimulus (2,3B) in the incubation vial. Mean ΔP/N value of the flies trained in this backward conditioning paradigm remained near zero, suggesting no memory, whereas the flies trained with the normal forward conditioning have significantly higher ΔP/N value than backward conditioning group (Figure 1D). This finding further supports the idea that the increment of ΔP/N for forwardly trained flies is exclusively attributable to the co-presentation of CS+ (attractive odor) and UCS (bitter/toxic food), which is the underlying principle behind the associative memory.

### The training paradigm induces protein synthesis dependent long-term aversive olfactory conditioning memory formation

Next, we set out to examine the longevity of the memory induced by the ‘attractive odor – bitter/toxic food’ coupling. For this purpose, 8-cycle trained flies were tested for their memory using the Y-maze assay after different time intervals. The results showed a steady decrement of memory performance (ΔP/N) of the trained flies over the course of 1 to 8 days after training (Figure 2A); memory is strongest when tested after 1 day, which gradually gets eliminated during the course of next 8 days. Till 7^th^ day memory is detectable (positive ΔP/N) whereas on 8^th^ day ΔP/N value is close to zero.

**Figure 2.**
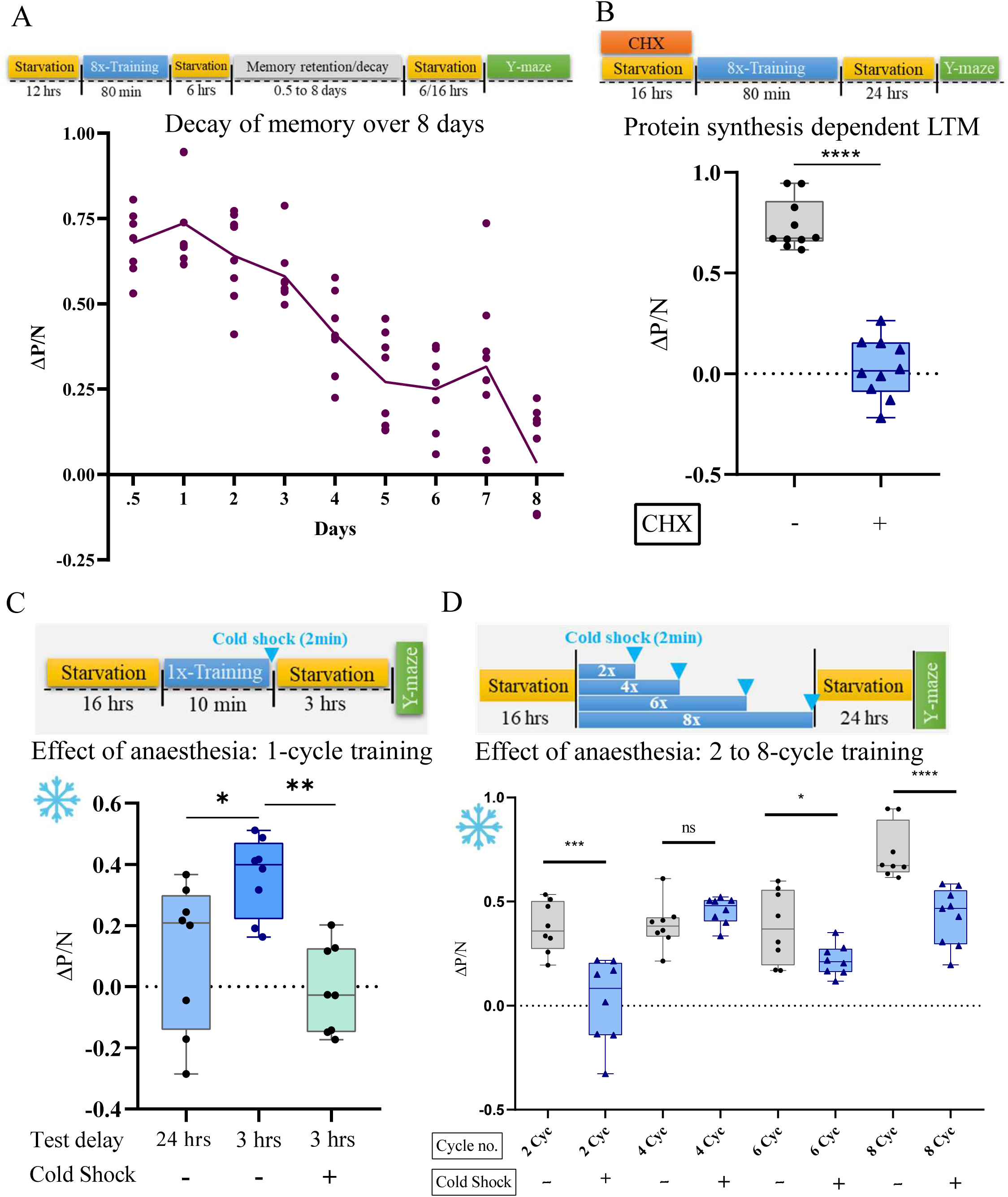
Deconstruction of the memory types induced by the “odor-bitter/toxic food” coupling associative memory. A. ΔP/N value of the 8 Cyc-trained flies were evaluated at different time intervals after training. The memory decay trajectory suggests memory retention for up to 7 days with gradual memory elimination over the course of 8 days. B. Inhibition of protein synthesis by CHX-feeding prior conditioning prevents the formation of aversive memory. In comparison, forward conditioning shows normal memory formation (same data points as Figure 1B). C. 1 Cyc trained flies fail to retain memory up to 24 hrs, but memory is present at 3 hrs after training, whereas cold-induced anesthesia eliminates this 3 hrs memory. D. ΔPI/N values of non-cold-shocked and cold-shocked groups are compared statistically. A sharp decrease in the value of ΔPI/N (near zero) in cold-shock group compared to ΔPI/N of non-cold-shocked group in 2 Cyc-trained flies indicates complete elimination of the 2 Cyc memory. In case of 4 Cyc-trained flies ΔPI/N remained unchanged after cold-shock; whereas for 6 Cyc and 8 Cyc trained flies, cold shock led to significant decrement in ΔPI/N but values were not near zero, indicating only a partial elimination of memory by cold induced anaesthesia (same data points as Figure 1C). N=8 biological replicates in each behavioral experiment (At least). Bars represent the Mean ± Standard Error of Mean (SEM). Unpaired two-tailed t test was performed for pair-wise comparision in all cases except 2C. Two-way ANOVA followed by Tukey’s multiple comparison test was performed for Figure 2C. **** signifies p value < 0.0001, *** signifies p value < 0.001, ** signifies p value < 0.01, * signifies p value < 0.05, “ns (non-significant)” signifies p value > 0.05.

As the memory elicited by 8 cycle training with the current paradigm lasts for over 7-8 days, it may be postulated as “long-term memory (LTM)”. Previous reports have established that long-term memory formation is dependent on the synthesis of new proteins to enable synaptic plasticity. Cycloheximide (CHX) is a protein synthesis inhibitor, which binds to the E-site of the 60S ribosome, thereby blocking the translocation step of translation (Schneider-Poetsch et al. 2010; Tully et al. 1994). To test whether the current paradigm induces a protein-synthesis dependent memory, protein synthesis was blocked prior to the memory induction. 35 mM CHX was added to the agar media and the flies were allowed to feed on it for 16 hrs immediately prior to conditioning protocol; this was followed by the training and assay of memory in Y-maze. CHX-fed CuSO_4_-trained flies showed significantly reduced ΔP/N value with respect to the flies fed with normal food (Figure 2B), i.e., formation of the 24-hr memory was severely affected by inhibition of new protein synthesis, proving that the ‘odor – bitter/toxic food’ associative memory formed by our paradigm is a protein synthesis dependent long-term memory (LTM).

In contrast, since 1 cycle training did not elicit a 24-hr LTM (Figure 1C), we checked whether it led to the formation of short-term memory (STM) instead. Y-maze assay performed 3 hours after 1 cycle training resulted in a positive ΔP/N (Figure 2C), indicating the formation of STM after 1 cycle.

### The aversive associative memory is a combination of anaesthesia resistant memory (ARM) and long-term memory (LTM)

Classically, long-lasting memory in *Drosophila* has been categorized into two different forms with clearly distinct molecular pathways: protein-synthesis dependent long-term memory (LTM) and anaesthesia resistant memory (ARM). As described previously for odor-shock conditioning, a brief cold shock is a useful tool to eliminate the *anaesthesia sensitive memory (ASM)* (Murakami et al. 2017; Noyes et al. 2020). To test whether the aversive memory induced by the olfactory-bitter/toxic food conditioning paradigm has a component of ARM, the trained flies were subjected to a cold shock by dipping the vials containing the flies into ice for 2 minutes immediately after training.

To analyse the relative contribution of ARM vs ASM in the aversive conditioning memory paradigm, effect of cold shock was tested on STM generated by a single training cycle (Figure 2C) and on the LTM induced by 2, 4, 6 and 8 training cycles (Figure 2D). When cold shock anaesthesia was applied to the single cycle trained flies, the 3-hr memory was found to be completely eliminated (Figure 2C), this indicates that the short-term memory (STM) formed through this paradigm is sensitive to anaesthesia and hence is AS-STM (anaesthesia sensitive short-term memory), and not ARM.

Subsequently, cold shock was applied on flies trained with higher number of training cycles. It was observed that the memory index (ΔP/N) of the 2-cycle trained cold-shocked flies was significantly reduced (near zero) compared to the 2 cycle non-cold-shocked flies after 24 hrs (Figure 2D). As the 24 hr memory elicited by 2-cycle training is eliminated by anaesthesia induction, the 2-cycle memory may be designated as anaesthesia sensitive LTM (AS-LTM).

In contrary, the 4-cycle trained group shows no loss of memory upon cold shock, hinting at the appearance of ARM by 4-cycle training. Interestingly, 6-cycle and 8-cycle trained flies showed significant reduction of memory upon cold shock, but the memory did not reach zero (Figure 2D). This demonstrates that the memory at 6 or 8 cycles are composite of AS-LTM and ARM; anaesthesia eliminates the ASM component, leaving the LTM component intact.

To further test whether the 2-cycle training generates a protein synthesis dependent LTM, cycloheximide (CHX, protein synthesis inhibitor) was added to the food and the flies were allowed to feed on this CHX-food for 16-hours prior to 2-cycle aversive conditioning and memory was tested after 24-hours. Surprisingly, this disrupts memory (Figure 3A), establishing 2-cycle memory as protein synthesis dependent LTM. It is known that generation of LTM requires post-training consolidation, and cold-anaesthesia presumably affects this consolidation, we sought to confirm that 2-cycle training is followed by memory consolidation. In the next experiment, cold shock was given 1 hour after 2 cycles of aversive training to spare the consolidation process and compared with the result of immediate cold shock. While immediate cold shock led to complete disruption of memory (near zero) the delayed cold shock only reduced the memory, but failed to eliminate the 24-hours memory fully (Figure 3B). This convincingly shows that 2-cycle training leads to protein-synthesis and consolidation dependent LTM.

**Figure 3.**
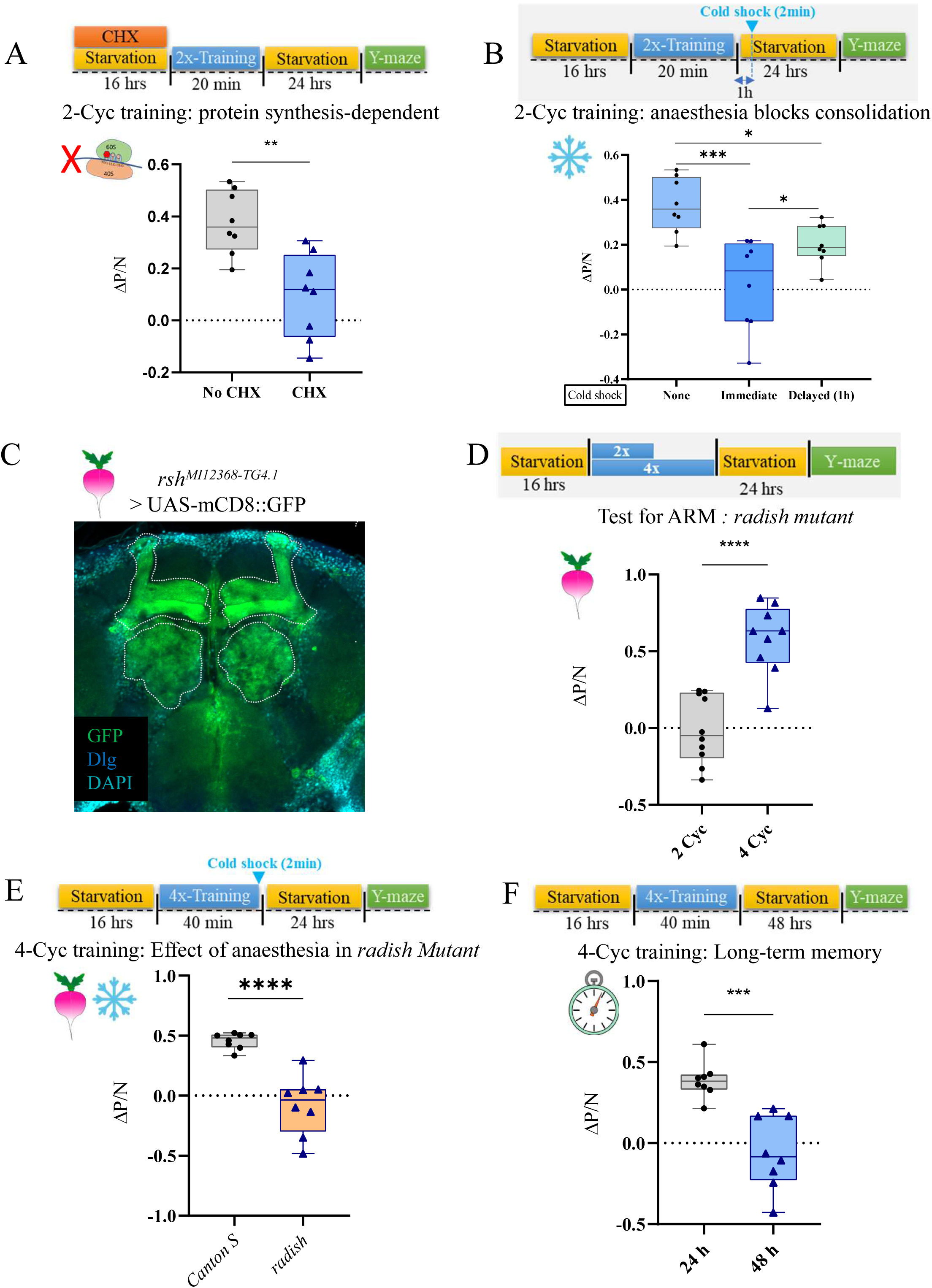
This aversive associative memory is a combination of both Anesthesia Resistant Memory (ARM) and Long-Term Memory (LTM) A. Feeding of protein synthesis blocker CHX prior conditioning eliminates 2 Cyc-trained 24 hrs aversive olfactory memory. B. Delayed cold shock, 1 hr after the 2 Cyc training in the *Canton S* (wild-type) flies, i.e., failed to annihilate the memory fully. C. *radish mutant (rsh^MI12368-TG4.1^)* line, containing GAL4 driven by the *radish* promoter, was crossed with *UAS-mCD8GFP* reporter line. GFP expression was observed throughout the different lobes of the Mushroom body and as well as within the antennal lobes. D. 2 Cyc trained *radish mutant* flies did not retain any traces of aversive olfactory memory after 24 hrs. In contrast, 4 Cyc-trained flies exhibited a greater positive ΔPI/N when compared to 2 Cyc trained flies. Hence, 4 Cyc training cycle induces a Radish-dependent ARM component. E. For the *radish mutant* flies, administering cold shock mediated anesthesia immediately after 4X training erased the 24 hrs memory. F. No memory traces were observed in the wild-type 4-cycle trained flies 48 hrs after training. N=8 biological replicates in each behavioral experiment (At least). Bars represent the Mean ± Standard Error of Mean (SEM). Two-way ANOVA followed by Tukey’s multiple comparison test was performed for Figure 3B. Unpaired two tailed t test was performed for other comparisons. **** signifies p value < 0.0001, * signifies p value < 0.05, “ns (non-significant)” signifies p value > 0.05.

In summary, 1-cycle and 2-cycle training only leads to AS-STM and AS-LTM respectively, whereas 4 or more cycles induces the ARM pathway in addition to the ASM. 6 or 8-cycle training elicits both ARM and AS-LTM.

### Role of *radish* gene on the ARM induced by the aversive conditioning

The molecular mechanism and pathway for the formation of Anaesthesia Resistant Memory have been described previously (Tully et al. 1994). *radish* gene, which codes for a Rap-like GTPase activating protein (RapGAP), has been found to be required exclusively for the ARM formation (Tully et al. 1994) and it is expressed throughout the different lobes of MB and antennal lobe (Figure 3C). Hence, to understand the contribution of ARM in the aversive memory, *radish* mutant flies were tested for their ability to form the aversive olfactory memory. It was observed that in *radish* mutant flies, 2-cycle memory was absent, indicating that the formation of 2-cycle memory is Radish-dependent (Figure 3D). So, combining the cold shock and radish mutant results, it may be stated that this 2-cycle memory is AS-LTM whose formation is dependent on Radish function.

In contrast, memory formation by 4 cycles of training was unperturbed in *radish* mutant flies (Figure 3D). This result, in combination with the cold-shock experimental result, where cold shock also failed to eliminate 4-cycle memory (Figure 2C), sufficiently establishes that the LTM induced by our paradigm is a combination of ARM and AS-LTM. To establish this further, 4-cycle training of *radish* mutant flies was followed by 2 min cold shock, and memory test after 24 hrs revealed no memory, presumably because lack of Radish blocked ARM component and cold-shock eliminated the AS-LTM component (Figure 3E). This unequivocally establishes that 4-cycle training is sufficient to elicit both ARM and AS-LTM in the fly brain. As ARM is known as mid-term memory lasting up to 24 to 48 hrs, we sought to check the longevity of 4-cycle training memory. Flies showed evidence of strong memory when Y-maze assay performed 24 hrs after training; but when the same was performed 48 hrs after training, no trace of memory was observed (Figure 3F). Hence, 4-cycle training induces a combination of ARM and AS-LTM, lasting for at least up to 24 hrs and decaying by 48 hrs after training.

In summary, it may be concluded that the aversive memory induced by the associative training coupling the bitter/toxic CuSO_4_ and attractive odorant 2,3B leads to the formation of strong long-term memory, which is a combination of short-term ASM, medium-term ARM and a long-term protein-synthesis dependent ASM. The anaesthesia-sensitive short-term memory (AS-STM) is induced by 1 cycle training, lasting for less than 24 hrs. Memory of 2-cycle training is protein synthesis dependent and lasts for up to 24 hrs, it is anaesthesia sensitive, hence may be called AS-LTM, and it also requires Radish protein for its formation. 4-cycle training induces an anaesthesia resistant memory (ARM) which also requires Radish, but lasts up to 24 hrs, decaying before 48 hrs (Figure 3F). Training of 6 or 8 cycles leads to protein synthesis dependent LTM (Figure 2B), which is only partially anaesthesia sensitive, lasting up to 7 days, presumably a composite of AS-LTM and ARM (Table 1).

**Table 1.**
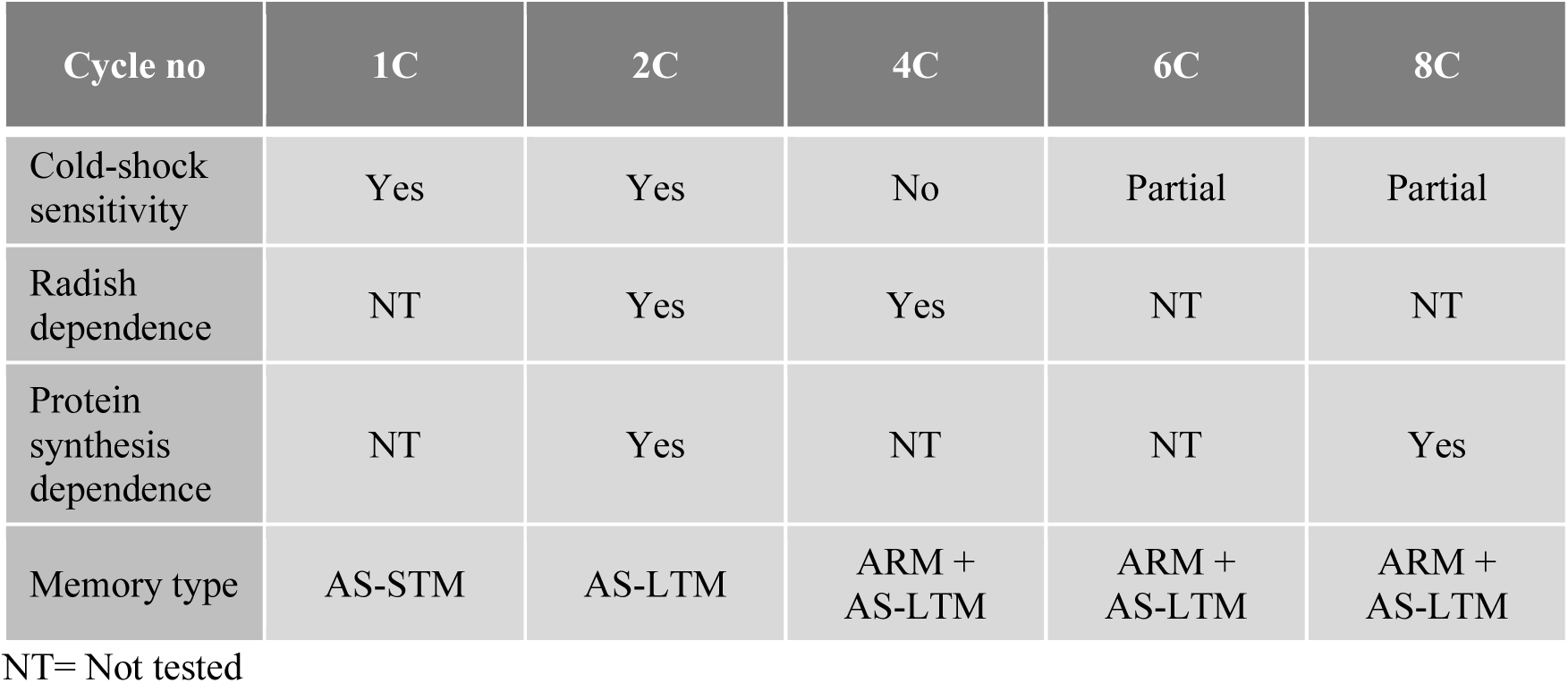
Summary of memory types formed in different cycles of the aversive memory paradigm.

| Cycle no | 1C | 2C | 4C | 6C | 8C |
| --- | --- | --- | --- | --- | --- |
| Cold-shock sensitivity | Yes | Yes | No | Partial | Partial |
| Radish dependence | NT | Yes | Yes | NT | NT |
| Protein synthesis dependence | NT | Yes | NT | NT | Yes |
| Memory type | AS-STM | AS-LTM | ARM + AS-LTM | ARM + AS-LTM | ARM + AS-LTM |

NT= Not tested

### Suppression of neurotransmission from Mushroom body neurons and PPL1 dopaminergic neurons during conditioning causes defect in LTM

It has been previously established that the olfactory associative memory (both appetitive and aversive) is mainly stored in the higher brain center neurons of the mushroom body (MB) in *Drosophila* (Aso et al. 2014). Immunofluorescence studies followed by confocal imaging revealed that, among other driver lines, *MB247-Gal4* line drives in the largest number of neurons (α, β, and γ lobes) of the MB Kenyon Cells (schematic in Figure 4A) with very little expression elsewhere in the brain (Pech et al. 2013) (Supplementary Figure S3A). Relying on the expression in the α, β and γ lobes of the MB, the *MB247-Gal4* line has been considered as an ideal driver line for checking whether this aversive conditioning is mushroom body dependent. To examine the role of MB in the aversive learning, the neurotransmission of mushroom body calyx neurons was suppressed during the conditioning experiment; a temperature-sensitive *shibire* gene (*shibire^ts^*) was ectopically expressed in the mushroom body neurons (*MB247-Gal4>UAS-shibire^ts^*) and neurotransmission from MB neurons were blocked by shifting to high temperature (30°C) 30 min prior to training followed by Y-maze assay at room temperature. ΔP/N value of *MB247-Gal4>UAS-shibire^ts^* flies is closer to zero, confirming LTM defect (Figure 4A). However, genotype controls: *MB247-GAL4*/+ and *UAS-shi^ts^*/+ exhibited significantly higher ΔP/N value with respect to *MB247-Gal4>UAS-shibire^ts^*, suggesting normal aversive LTM formation (Figure 4A). Moreover, flies carrying the experimental genotype *MB247-Gal4>UAS-shibire^ts^* when trained at 18℃ (no induction of *shibire^ts^*), ΔP/N value is significantly higher from experimental genotype at 30℃, indicating LTM formation remained intact (Figure 4A).

**Figure 4.**
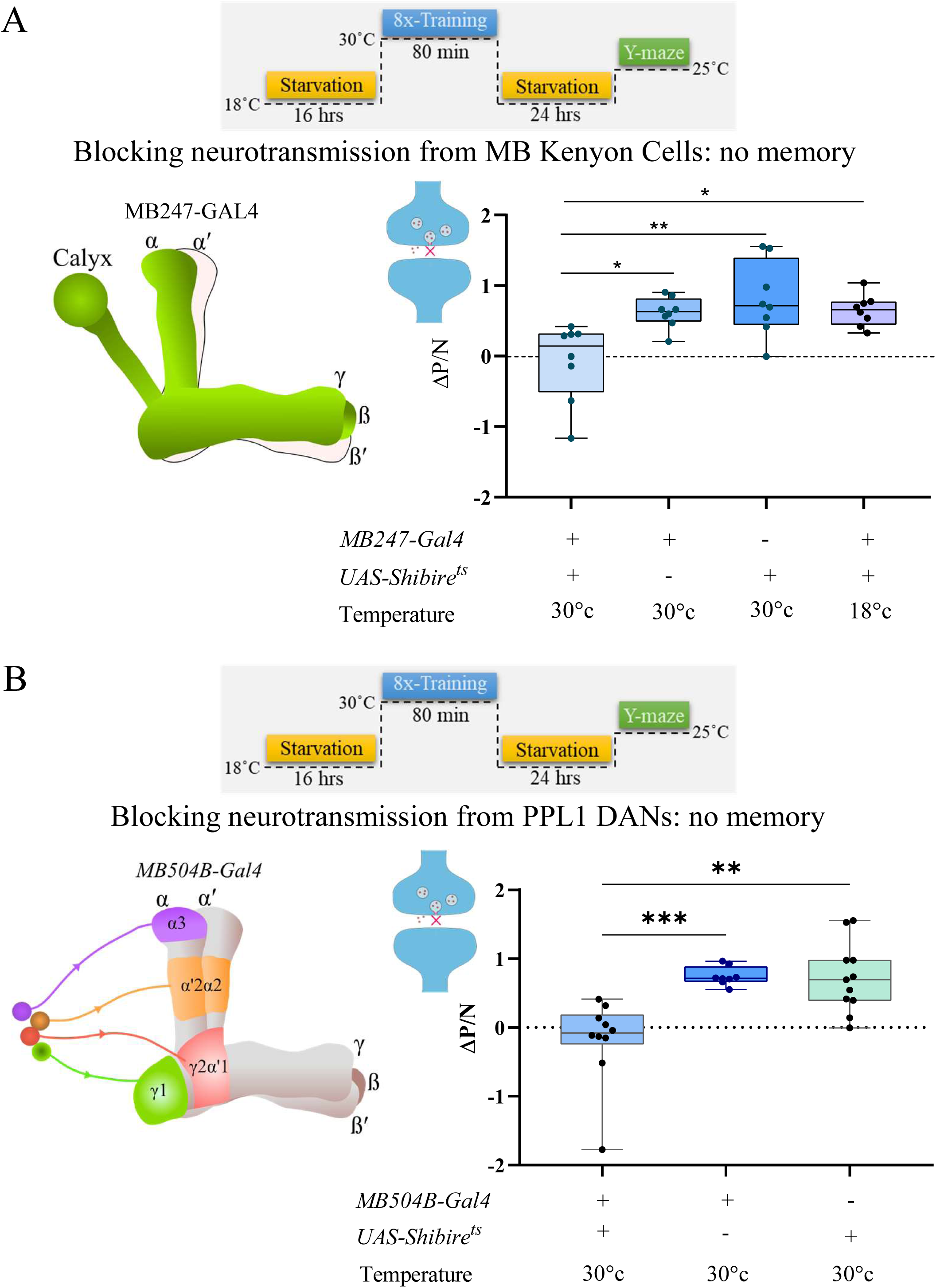
Suppression of neurotransmission from MB Kenyon cell neurons and PPL1 neurons during conditioning leads to LTM defect. A. Neurotransmission of mushroom body Kenyon cell neurons was selectively inhibited during the conditioning experiment by ectopically expressing a temperature-sensitive *Shibire* gene (*shibire^ts^*) in the *MB247-Gal4* positive neurons and elevating the temperature to 30°C during training. ΔPI/N of *MB247-Gal4>UAS-shibire^ts1^* flies exhibited nearby zero value, confirming LTM defect. Genotype-control flies carrying only *MB247-GAL4* (*MB247Gal4/+)* and only *UAS Shibire^ts1^* (*UAS Shibire^ts1^/+)* trained at 30°C showed significantly higher ΔPI/N than *MB247-Gal4>UAS-shibire^ts1^* flies, confirming intact normal aversive LTM. When the flies of the genotype *MB247-Gal4>UAS-shibire^ts1^* were trained at 18℃, the trained flies exhibited a significant increase in ΔPI/N than the *MB247-Gal4>UAS-shibire^ts1^* flies trained at 30℃; indicating proper induction of memory. B. When the neurotransmission from PPL1 neurons was silenced (*MB504B-Gal4>UAS-Shibire^ts1^*) during conditioning, ΔPI/N values remain nearby zero, indicating LTM defect. LTM formation remains unaffected in genotype control (*MB504BGal4/+* and *UAS Shibire^ts1^/+*) with the same temperature regime. N=8 biological replicates in each behavioral experiment (At least). Bars represent the Mean ± Standard Error of Mean (SEM). Two-way ANOVA followed by Tukey’s multiple comparison test was performed for Figure 4A. One way ANOVA followed by Tukey’s multiple comparison test was performed for Figure 4B. *** signifies p value < 0.001, ** signifies p value < 0.01, “ns (non-significant)” signifies p value > 0.05.

From previous studies it is known that aversive memory induced by odor-shock association, is contributed by a special set of dopaminergic neurons (DANs) which form synaptic connections with MB Kenyon cells (Claridge-Chang et al. 2009). Two separate clusters of dopaminergic neurons also project their axons to different lobes of MB, enabling the transmission of either reward or punishment signal to specific regions of MB mediating associative learning. In case of aversive learning, posterior DANs of PPL1 cluster relay negative reinforcement onto the MB neurons (Cognigni et al. 2018). *MB504B-Gal4* selectively labels different clusters of PPL1 neurons (schematic in Figure 4B Supplementary Figure S3B). We used the *MB504B-Gal4* driver line to drive *UAS-shibire^ts^* transgene in these PPL1 neurons and the neurotransmission of these PPL1 neurons to MB lobes was silenced by temperature induction (30°C) during training. LTM formation was found to be abolished in flies where PPL1 neurotransmission was blocked by elevating temperature to 30°C in flies of the genotype *MB504B-Gal4*>*UAS-shibire^ts^*. LTM formation remains unaltered in genotype control flies, *MB504B-Gal4/+* and *UAS-shi^ts^*/+ (Figure 4B).

### Aversive conditioning leads to Ca^2+^ surge in MB γ-neurons and synaptic elaboration

Formation of long-term memory involves calcium mediated signaling to the nucleus leading to expression of genes mediating synaptic plasticity, which in turn, is a signature of LTM. To identify if our training regime induces an intracellular calcium signal, we used a bipartite **<u>T</u>**ranscriptional **<u>R</u>**eporter of **I**ntracellular **<u>C</u>**a^2+^ (TRIC) reporter (Gao et al. 2015) driven in the mushroom body neurons. TRIC is composed of two fusion proteins: calmodulin is fused with transcription activation domain (CaM-AD) and CaM target peptide is fused with GAL4 DNA binding domain (CaM target-DBD). Enhanced calcium causes structural change in the CaM protein, bringing the fused AD in close proximity with the DBD, triggering GFP expression downstream to UAS sequence; a constitutively expressed RFP works as an internal reference for control. The *MB247-Gal4>UAS-TRIC* flies were trained with aversive conditioning paradigm followed by dissection and immunolabeling of their brains and quantification of GFP and RFP from the γ-lobes of the MB (Supplementary Figure S4A and schematic in Figure 5A). We observe a significant enhancement in the GFP/RFP ratio in the trained fly (Figure 5B), indicating an enhanced neuronal activity indicated by enhanced intracellular calcium in the γ neurons of the trained flies. In order to check whether the enhanced calcium culminates in synaptic plasticity, we immunolabeled and quantified a post-synaptic marker Discs-large (Dlg) from the same γ-lobes of the MB. Significantly enhanced level of Dlg was observed in the trained flies indicating an elaboration of synaptic arborization in the mushroom body neuropil region (Figure 5C); this demonstrates synaptic plasticity is induced by the aversive conditioning paradigm. Also, no synaptic arborization was observed for backwardly trained flies (Supplementary Figure S4B).

**Figure 5.**
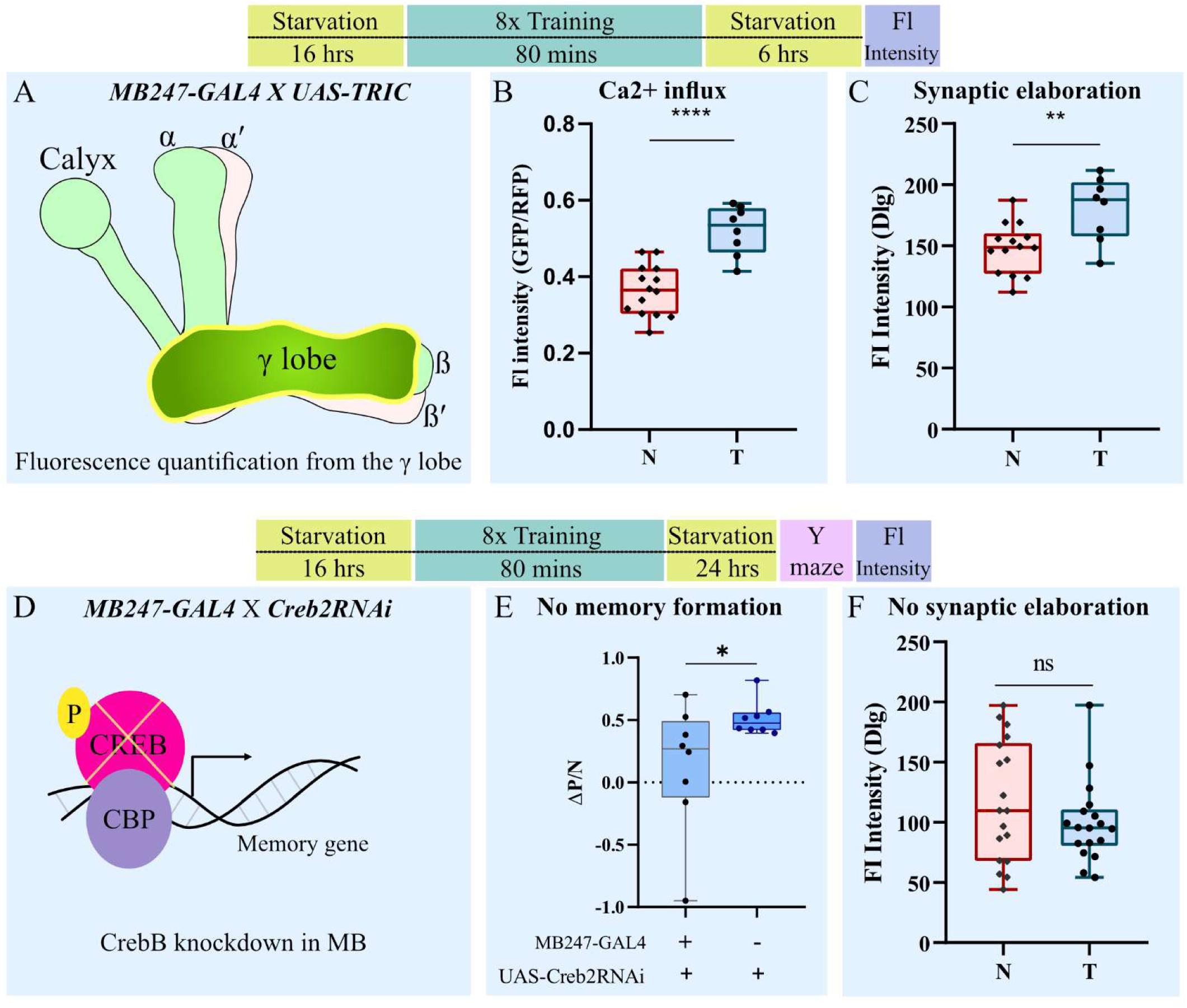
The aversive LTM is associated with Ca^2+^ surge and synaptic elaboration in the MB γ-neurons and requires CrebB-mediated transcription. A. Schematic diagram of different lobes of the mushroom body of adult fly brain. Fluorescence quantification was performed from the γ lobes of the mushroom body. B. Fluorescence quantification from mushroom body γ lobes of adult fly brains of the genotype *MB247>UAS TRIC* labelled with anti-GFP (green) and anti-RFP (red). Elevated ratio of GFP/RFP intensity in the MB neurons indicates enhanced Ca^2+^ surge in the trained flies. C. Quantification of pre-synaptic protein Dlg in the mushroom body γ neurons by immunofluorescence quantification (anti-Dlg) of adult fly brains shows trained flies have significantly higher Dlg than the that of the naïve flies, suggested enhanced synaptic plasticity in MB γ-neurons. D. Knockdown of CrebB by driving the expression of an RNAi in the mushroom body neurons by the *MB247-Gal4* driver line E. Knockdown of CrebB caused aversive olfactory memory defect. F. Quantification of Dlg-immunofluorescence intensity in CrebB-knocknown mushroom body shows no significant increase in trained flies compared to that of the naïve flies. N=8 biological replicates in each experiment (At least). Bars represent the Mean ± Standard Error of Mean (SEM). Unpaired two-tailed t-test was performed for each case. **** signifies p value < 0.0001, ** signifies p value < 0.01.

### CrebB mediated transcription is required for the long-term aversive conditioning memory

CrebB (cyclic-AMP response element binding protein) is a transcription factor which binds with the CRE (cyclic-AMP Response Element) region upon receiving neuronal activity dependent calcium signal during learning and memory formation. Many memory-related genes harbour CRE sequences in their promoters. Binding of CrebB to these sites enables recruitment of the histone acetyltransferase CBP, which acetylates chromatin to support activity-induced transcription necessary for synaptic plasticity and long-term memory formation (Figure 5D). Hence, to check if CrebB is involved in the aversive LTM, we performed RNAi-mediated knockdown of *CrebB* (supplementary Figure S4C) in the MB neurons during aversive conditioning and observed LTM to be defective (Figure 5E), whereas, flies containing only *UAS-CrebB RNAi* showed normal memory; this entails the requirement of CrebB-mediated transcription for aversive conditioning memory formation. Further to corelate the behavioral performance with probable synaptic plasticity in these flies, the brains of the *MB247-Gal4>UAS-CrebB RNAi* flies were dissected 6 hrs after conditioning, immunolabelled against post-synaptic protein Discs large (Dlg) and imaged in confocal microscope; levels of Dlg was measured by fluorescence quantification from the MB γ lobes (Figure 5F). Non-significant FI intensity of Dlg is observed between naïve and trained flies (Figure 6G), suggesting no synaptic elaboration in MB neurons upon *CrebB* knockdown. Also, no synaptic elaboration has been observed for the wild-type backward trained flies. These results establish that the synaptic elaboration (plasticity) in the MB γ lobes is induced by conditioning and is dependent of CrebB function.

**Figure 6.**
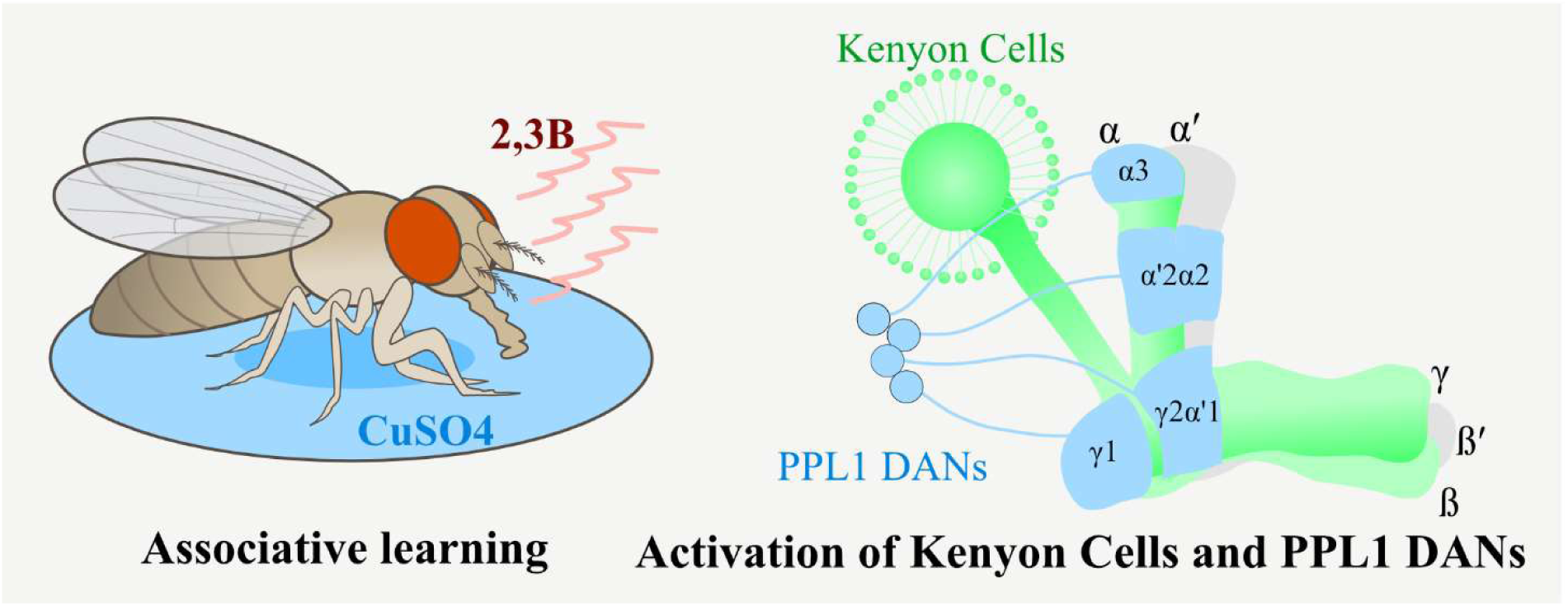
Schematic showing the neuroanatomical correlates of the Drosophila odor-CuSO_4_ associative memory formation. Traumatic ‘odor-stress’ conditioning memory formation requires neurotransmission from MB Kenyon Cells and PPL1 DANs.

## Discussion

Long-lasting memory formation relies on robust synaptic plasticity and orchestrated molecular mechanisms. Memories of traumatic experiences trigger a stressful memory which sustains for long duration and often lead to post-traumatic stress disorder (PTSD). Understanding the unique fundamental features of this memory would be useful to device strategies to overcome the problems associated with it. Our novel aversive odor-stress conditioning memory paradigm in *Drosophila*, pairing a single attractive odorant with a sustained stress stimulus, induces a composite memory comprising both anesthesia-resistant memory (ARM) and consolidation-dependent long-term memory (LTM). This paradigm offers distinct advantages over existing gustatory or olfactory memory assays and reveals new insights into multimodal memory encoding in the fly brain. One significant feature is the extended persistence of the memory for up to seven days, surpassing durations typically observed in gustatory paradigms using either appetitive (e.g., sucrose) (Tully et al. 1994; Isabel et al. 2004) or aversive (e.g., quinine) (El-Keredy et al. 2012) cues as conditioned stimuli. The use of a single odorant as conditioned stimulus (CS), instead of the conventional two-odor training simplifies the training method and makes it possible to potentially detect training-induced valence-reversal of the odorant (observed in case of larval training by slightly modified paradigm, Majumder et al, unpublished observation).

One possible caveat of this novel paradigm may be the possibility of generation of a parallel appetitive memory during aversive conditioning: In this experiment, the trained flies were given Agar-Sucrose-CuSO_4_ food whereas the naïve group was given Agar-Sucrose food. Sucrose was added in both vials to ensure ingestion of the food, but being a known appetitive reinforcer, it may also elicit an appetitive learning by associating the sweet taste of sucrose with the attractive odor 2,3B, although this concurrent appetitive learning is also reported to be induced the odor-shock associative memory (Felsenberg et al. 2018).

Our results highlight a cycle-dependent, parallel and independent emergence of STM, ARM, and LTM. Fewer training cycles induce ARM, traditionally characterised as protein synthesis-independent and Radish-dependent and higher number of cycles induce a robust and protein synthesis–dependent LTM. Blocking ARM by *radish* mutation does not impact anaesthesia-sensitive LTM generation and blocking ASM by cold shock does not affect ARM. Moreover, cold shock in *radish* mutant background effectively abolishes the entire memory, confirming independent and simultaneous induction of both types of memory. To summarize, our finding establishes that the emergence of these three types of memories happens through parallel, presumably by the action of independent molecular pathways, confirming previous findings in other paradigms.

Our findings establish the requirement of inputs from the mushroom body (MB) Kenyon cells and the dopaminergic PPL1 neurons during training (Figure 6). Moreover, increased calcium signalling in γ-lobe MB neurons and synaptic elaboration suggest that structural changes in synapses guides behavioural plasticity underlying long-term memory storage.

In sum, our odor-stress conditioning paradigm induces a long-lasting, multimodal memory that involves simultaneous or parallel generation ARM and LTM components, requiring the classical memory circuitry, and mediated by calcium induced synaptic plasticity. The simplicity of the training paradigm and multimodality & longevity of the memory make it a powerful platform for deciphering classical questions in memory formation, consolidation, retention and their anomalies in human neurological disorders.

## Declaration

The authors declare no competing interest.

## Acknowledgements

AD laboratory received research grants from Govt. of India: Science and Technology Research Board-DST (ECR/2017/002963), Department of Biotechnology Ramalingaswami fellowship (BT/RLF/Re-entry/11/2016) and Department of Biotechnology (BT/PR53805/BMS/85/101/2024); from Govt. of West Bengal: WBDSTBT Core Research Grant (2297(SANC.)/STBT-13015/10/2024); and intramural research grant from IIT Kharagpur, India. We acknowledge the Confocal Microscope purchased under DST-FIST grant conferred on the Department of Bioscience and Biotechnology (erstwhile School of Bioscience), File no. SR/FST/LS-I/2019/595(C). SM is recipient of Prime Minister Research Fellowship (PMRF). The GD lab is funded by extramural funds from the Pratiksha Trust Extra-Mural Support for Transformational Aging Brain Research grant (EMSTAR/2023/SL03) and the Department of Biotechnology, Government of India grant (BT/PR51490/MED/ 22/361/2024). The lab also receives intramural funding support from BRIC-NCCS, Pune.

## List of Acronyms

ΔPI: Differences in performance indices
1X to 8X: 1 cycle to 8 cycle
2,3 BD: 2,3 Butanedione
AD: Activation Domain
ARM: Anaesthesia Resistant Memory
ASM: Anaesthesia Sensitive Memory
Brp: Bruchpilot
BSA: Bovine Serum Albumin
CaM: calmodulin
cAMP: cyclic adenosine monophosphate
cDNA: complementary DNA
CHX: Cycloheximide
CREB: cAMP response element-binding protein
CS: Conditioned Stimulus
CuSO_4_: Copper (II) sulfate
DAN: Dopaminergic neuron
DBD: DNA Binding Domain
Dlg: Discs-large
FI: Fluorescence Intensity
GAL4: galactose-responsive transcription factor
GFP: Green Fluorescent Protein
KC: Kenyon cell
LD: light/dark cycle
LTM: Long-term memory
MB: Mushroom Body
MBON: Mushroom body output neuron
MCH: 4-Methylcyclohexanol
mRNA: Messenger RNA
MTM: Middle-term memory
PAM: Protocerebral anterior medial
PBS: Phosphate Buffer Saline
PBT: PBS containing 0.1 % Triton X-100
PCR: Polymerase Chain Reaction
PI: Performance index
PKA: cAMP-dependent protein kinase A
PPL1: Protocerebral Posterior Lateral 1
PSD: Protein Synthesis Dependent
PSI: Protein Synthesis Independent
PTSD: post-traumatic stress disorder
RAP: Ras-related protein
RFP: Red Fluorescent Protein
RNAi: RNA interference
SEM: Standard Error of Mean
shRNA: Short hairpin RNA
STM: Short-term memory
STRV: Starvation
TRIC: Transcriptional Reporter of Intracellular Ca^2+^
TRN: Training
UAS: Upstream Activating Sequence
UCS: Unconditioned Stimulus

## Supplementary Figures

**Supplementary Figure S1.**
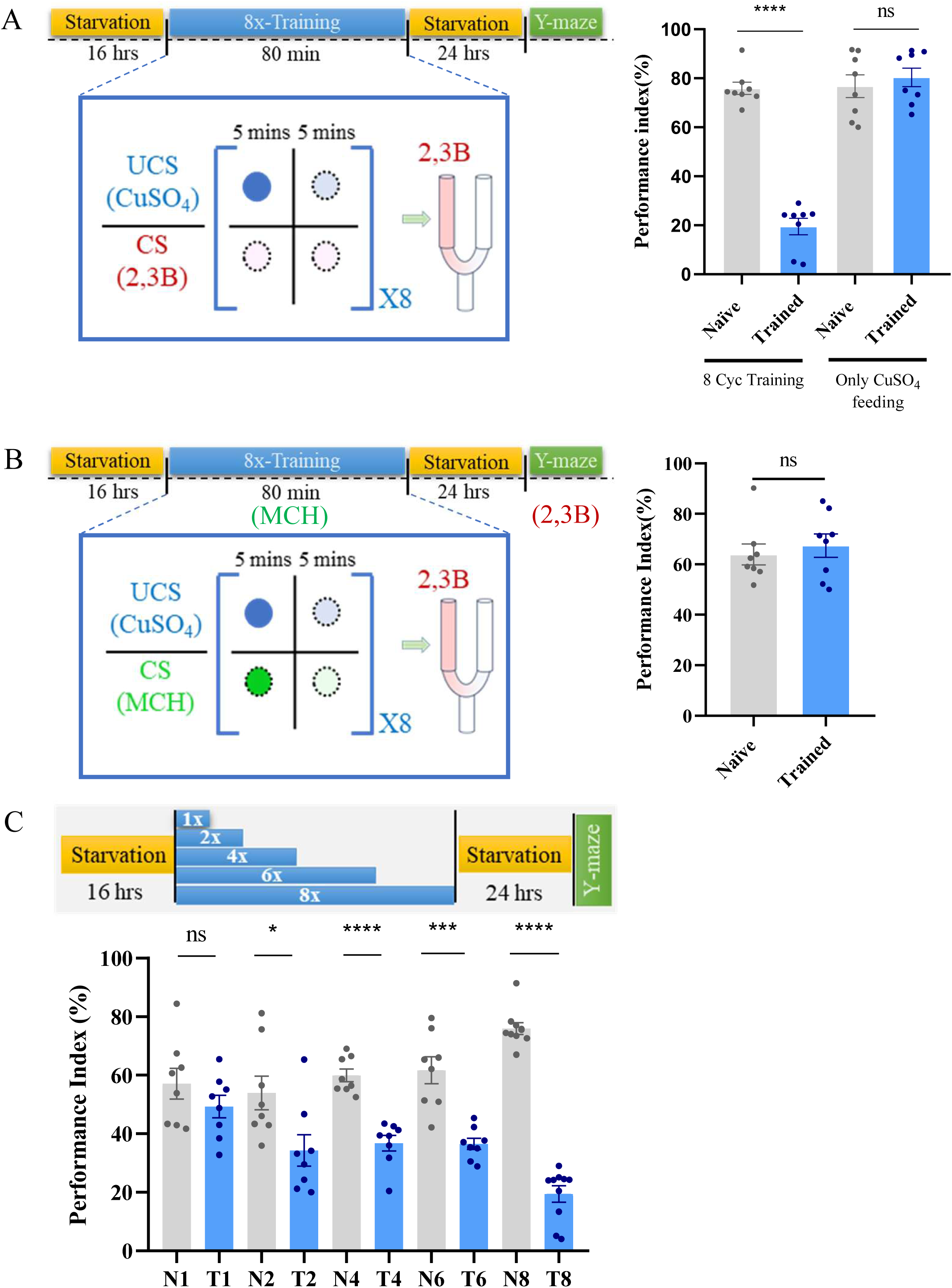
The formation of Aversive LTM arises specifically due to associative pairing between the odor stimulus and bitter/toxic food. A. Conditioning with Copper sulphate (CuSO_4_) containing food alone does not lead to decreased PI compared to naïve flies, i.e., no decreased attraction, towards the testing odorant during the odor choice assay, indicating unchanged innate odor response of the flies. B. Training the flies with a different odorant (MCH) and testing them with 2,3B does not lead to reduced PI; i.e., no change in odor bias towards 2,3B. C. Wild-type flies were trained with variable number of training cycles (1 Cyc, 2 Cyc, 4 Cyc, 6 Cyc and 8 Cyc) and their 24 hrs memory was assessed. Flies trained with one training cycle (1 Cyc) did not show any significant reduction in performance index with reference to the naïve controls. 2 Cyc-trained flies exhibited a significant drop in PI compared to naïve flies, indicating training induced memory formation. The 4 Cyc, 6 Cyc, and 8 Cyc) trained flies also showed a similar reduced attraction towards the trained odorant after 24 hrs. N=8 biological replicates in each behavioral experiment. Unpaired two-tailed t test was performed in each cases. Bars represent Mean ± Standard Error of Mean (SEM). **** signifies p value < 0.0001, *** signifies p value < 0.001, ** signifies p value < 0.01, * signifies p value < 0.05, ns (non-significant) signifies p value > 0.05.

**Supplementary Figure S2.**
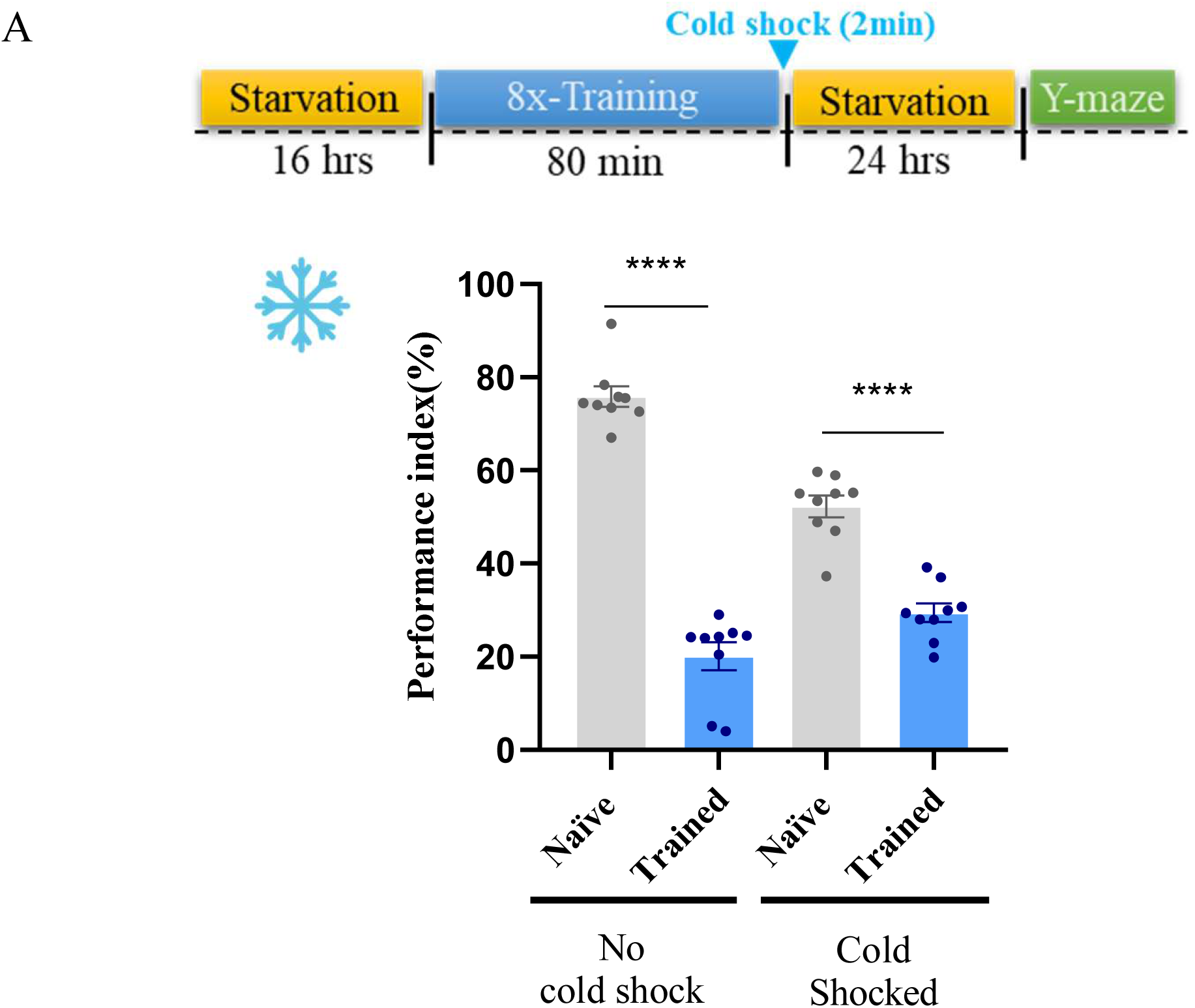
Aversive training for one cycle leads to the formation of anaesthesia-sensitive short-term memory. A. Post-training cold shock to 8X-trained flies does not lead to elimination of the aversive memory completely. N=8 biological replicates in each behavioral experiment (At least). Bars represent the Mean ± Standard Error of Mean (SEM). Unpaired two tailed t test was performed for Figure S2B. **** signifies p value < 0.0001, ** signifies p value < 0.01, * signifies p value < 0.05.

**Supplementary Figure S3.**
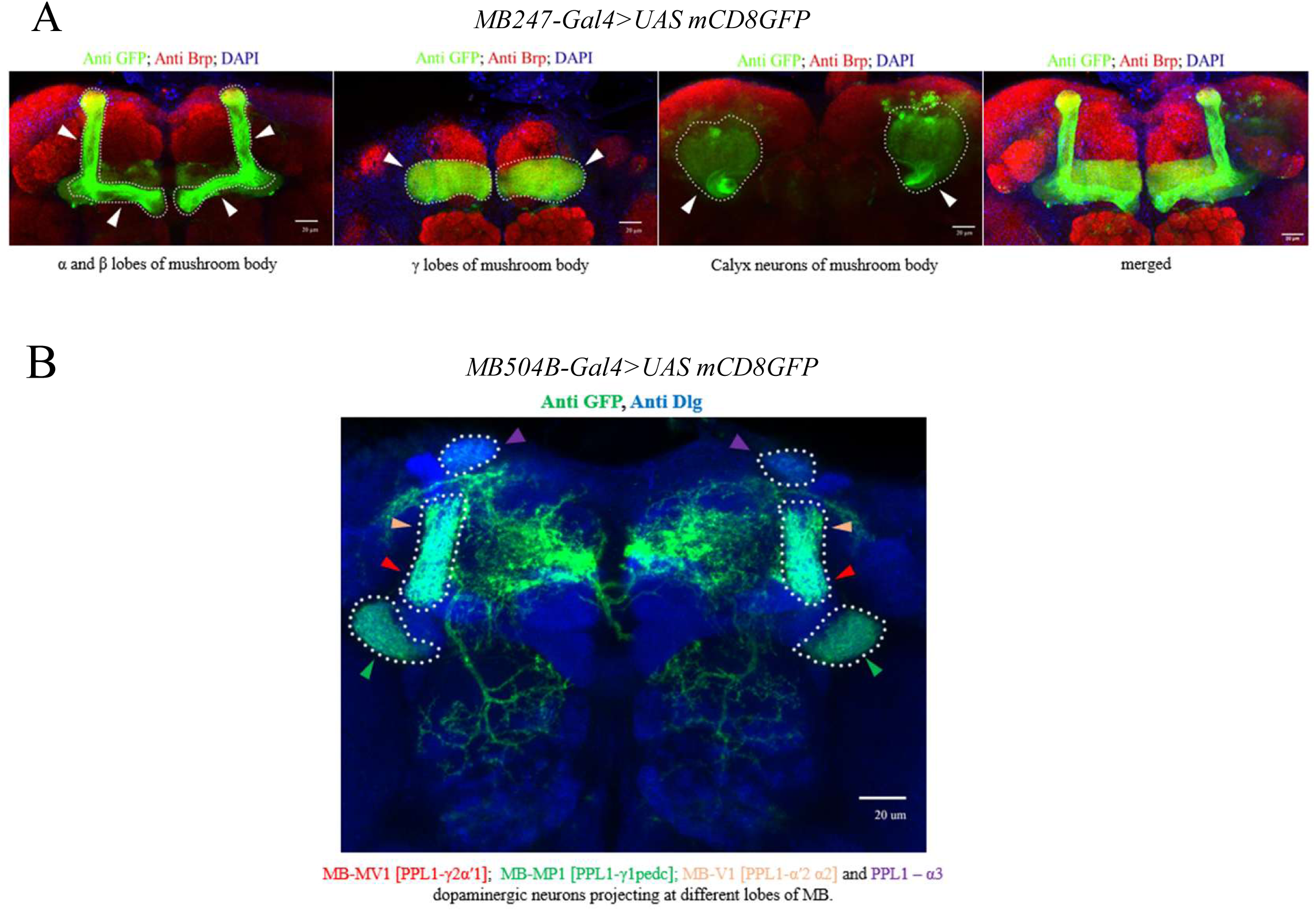
Expression patterns of the GAL4 lines used in the behavioral experiments. A. GFP expression in the α, β, and γ lobes along with calyx neurons of the mushroom body in *MB247 Gal4>UASmCD8GFP* fly line. B. GFP expression driven by MB504-GAL4 in the MB-MV1 [PPL1-γ2α′1] neurons, MB-MP1 [PPL1-γ1pedc] neurons, MB-V1 [PPL1-α′2 α2] neurons and PPL1-α3 dopaminergic neurons projecting at different lobes of MB.

**Supplementary Figure S4.**
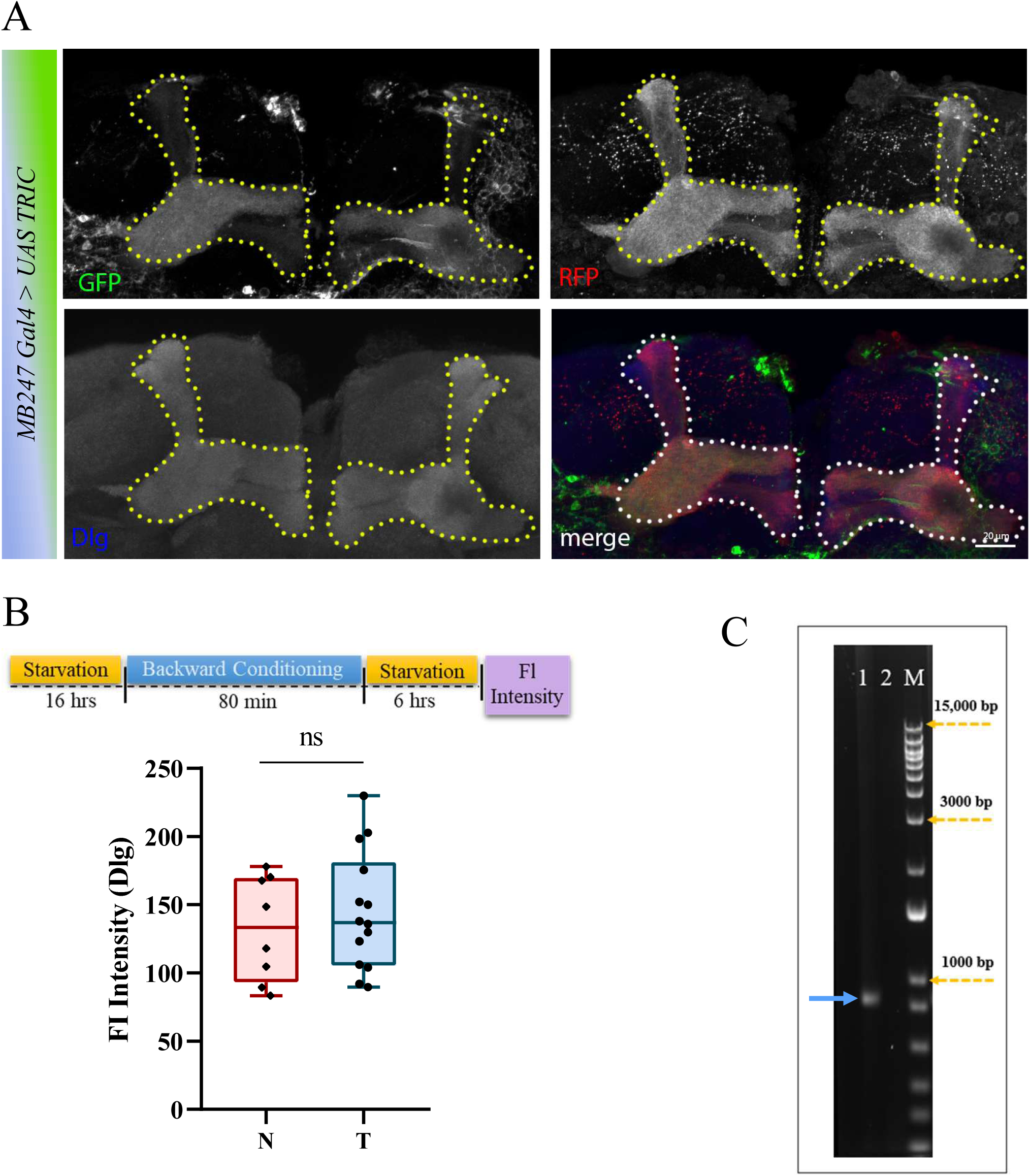
Validation of TRIC reporter expression in the MB neurons and efficacy of the CrebB-RNAi line. A. Adult fly brains of the genotype *MB247>UAS TRIC* labelled with anti-GFP (green), anti-RFP (red) and anti-Dlg (Blue) showing the regions from which fluorescence intensity (FI) quantification for Dlg, GFP, and RFP were performed in Figure 6B and 6C). B. Wild-type flies does not exhibit any synaptic elaboration upon backward conditioning. C. A PCR-based semi-quantitative method was used to validate the *UAS CrebB-RNAi* line. PCR amplified CrebB-cDNA isolated from the control and CrebB-knockdown heads were run in agarose gel. The CrebB-specific band was observed for the genotype control *Tub-GAL4/+* (blue arrow in lane 1). Corresponding CrebB-specific band was absent in the CrebB knockdown brain samples (*Tub-GAL4>UAS CrebB-RNAi* fly heads, lane 2), validating an efficient knockdown of CrebB by the RNAi. N=8 biological replicates in each behavioral experiment (At least). Bars represent the Mean ± Standard Error of Mean (SEM). Unpaired two-tailed t-test was performed for each case. ns (non-significant) signifies p value > 0.05.

